# Conservation of sensory pathways implies a localised change in the mushroom bodies is associated with cognitive evolution in *Heliconius butterflies*

**DOI:** 10.1101/2025.07.08.663516

**Authors:** Elizabeth A. Hodge, Denise D. Dell’Aglio, Antoine Couto, W. Owen McMillan, Max S. Farnworth, Stephen H. Montgomery

**Affiliations:** Evolution of Brains and Behaviour lab, School of Biological Sciences, University of Bristol, Bristol, United Kingdom; Smithsonian Tropical Research Institute, Gamboa, Panama; Evolution Genomes Behavior and Ecology, IDEEV, Université Paris-Saclay, Gif-sur-Yvette, France

**Keywords:** brain evolution, comparative neuroanatomy, compound eyes, neural circuits, Heliconiini, visual acuity

## Abstract

Acquisition of novel behaviour is reflected in changes in sensory investment or integration, but the exact nature of these changes is often unclear. Within the Neotropical butterfly tribe, Heliconiini, the genus *Heliconius* possess 4-fold larger mushroom bodies, an insect learning and memory centre, than closely related Heliconiini. Mushroom body expansion in *Heliconius* co-occurred with a dietary innovation, and is associated with systematic spatial foraging and extended lifespans. Heliconiini therefore offer an attractive system for studying how behavioural evolution is facilitated by changes in neural systems. *Heliconius*’ foraging relies on visual scene memories and, indeed, *Heliconius* have more stable visual long-term memory, and evidence of visual specialisation in the mushroom bodies. Here, we explore how vision-specific neuroanatomical and behavioural enhancement in *Heliconius* impacts sensory pathways upstream of the mushroom bodies by assessing investment across the eyes, sensory structures and projection pathways. Despite evidence of refinement in visually-based behaviour, we found no increased investment in visual structures, brain areas or pathways. This suggests that the rapid expansion of the *Heliconius* mushroom body occurred in a context of conserved detection and processing of visual cues, and that a localised shift within integrative brain centres facilitated the evolution of *Heliconius’* novel behaviours.

## Introduction

The sensory and cognitive abilities of a species are dependent on the neural systems that produce them. These systems can evolve through changes in cell number, including by replication of whole neural circuits and associated cell populations, or through altered patterns of connectivity, dendritic branching or neuromodulation (1,2). Evolutionary specialisations in neural systems have allowed species to exploit a diverse range of ecological niches, and thus, underpin the diversity of animal behaviour (2). The close correspondence between neural investment and ecological need is illustrated by divergent investment in the peripheral and central sensory processing structures of sensory stimuli across species (3–5). Consequently, shifts in behaviour can often be accompanied by corresponding shifts in sensory perception and processing. For example, changes in the peripheral structures, such as differences in the size or structure of the eyes, or in downstream visual neuropils, are often found between nocturnal and diurnal species (6). However, behavioural adaptations can also occur in the context of conserved sensory perception but altered integration of sensory responses in integrative brain centres (7–10). Understanding the extent to which peripheral and integrative systems can evolve independently, and which may most frequently underpin behavioural novelty, is a central challenge in evolutionary neurobiology (2).

The Neotropical butterfly genus *Heliconius* provides a suitable case study in adaptive evolution of brain centres associated with behavioural novelty. *Heliconius* have a unique dietary adaptation, which involves active pollen feeding, a foraging behaviour not observed in other butterflies (11–13). The innovation of pollen feeding has had profound implications for *Heliconius’* adult life history. Protein acquired through pollen feeding is linked to an extended lifespan and increased fecundity in *Heliconius* in comparison to their Heliconiini relatives (14–18), but with seemingly conserved juvenile development (19). Pollen feeding, therefore, has major fitness benefits, and *Heliconius* display derived foraging behaviours to support this behaviour (11,17,20). *Heliconius* collect pollen from a restricted range of floral plant resources in a systematic manner, repeatedly visiting specific plants in a spatially and temporally faithful manner, reminiscent of trap-line foraging in some Hymenoptera (11,21).

This systematic foraging behaviour relies heavily on spatial memory, likely through learning visual scenes to support recall of spatial routes (22–27). Therefore, *Heliconius* are thought to make greater use of spatial and temporal cues, and retain visual memories for longer periods of time than their non-pollen feeding close relatives in Heliconiini. Indeed, in Heliconiini there are modality-dependent shifts in memory stability between species, with *Heliconius* exhibiting greater retention of visual, but not olfactory, memories compared to outgroup genera (27,28).

These behavioural differences in foraging, learning, and memory have been strongly linked to the expansion of the mushroom bodies, the site of learning and memory in insects (29). In comparison to their close relatives in the Heliconiini tribe, *Heliconius* have ∼4-fold larger mushroom bodies relative to the rest of the brain, with around eight times the number of Kenyon cells, the intrinsic mushroom body neurons (30). Mushroom body expansion is also associated with internal specialisation, with an increase in the size and proportion of the mushroom body calyx receiving afferent visual input in *Heliconius* (30), as well as disproportionate expansion of specific mushroom body lobes, which make efferent connections to other brain regions (31). This major shift in mushroom body size and structure has occurred among relatively closely related species (*Heliconius* originated ∼8-14 mya; 32). Heliconiini, therefore, provide a tractable opportunity to study the links between the evolution of cognitive enhancement and neuroanatomical adaptations between species which, aside from the evolution of pollen feeding, are largely similar in terms of ecology, behaviour, and life history (13,33).

Until now, however, whether the evolution of *Heliconius*’ systematic foraging behaviour and mushroom body expansion was also accompanied by major changes in the upstream sensory organs and associated sensory neuropils was unclear. Given the reliance of *Heliconius* on visually guided foraging, it is a plausible hypothesis that adaptations in the visual pathway could enable *Heliconius* to utilise visual cues in ways their non-pollen feeding outgroup Heliconiini relatives cannot. Mushroom body expansion in *Heliconius* could, therefore, have occurred in response to increased visual capacities, rather than a direct response to selection linked to the information storage requirements of spatial foraging. To determine the extent to which changes in the integrative circuits of the mushroom bodies and downstream targets facilitate *Heliconius’* cognitively demanding foraging behaviours, variation in the sensory structures peripheral to the mushroom bodies must be considered.

In insects, variation in ommatidia of the compound eye are responsible for many of the key components of visual perception, including the ability to distinguish between different spectral wavelengths, different light intensities, different degrees of detail and temporal resolutions (34–37). Like other insects, visual information in Heliconiini is relayed from the eye through a series of visual neuropils making up the optic lobe (lamina, medulla, accessory medulla, lobula, ventral lobula and lobula plate in Heliconiini) before their output is relayed to integration centres of the brain through distinct populations of sensory projection neurons (38–40). Similarly, olfactory input is received from the antennae and processed initially in the glomeruli of the antennal lobe, the primary olfactory neuropil, before being relayed to other brain centres, primarily the mushroom bodies and lateral horn (41,42), by olfactory projection neurons. In previous work, volumetric investment in the medulla, as the largest visual neuropil, and the ventral lobula, as a likely relay point to the mushroom body, was assessed across Heliconiini, with no evidence of divergent evolution in *Heliconius* (30), but major aspects of the sensory pathways remain unassessed.

Here, we explore how sensory systems upstream of the mushroom bodies vary between *Heliconius* and outgroup genera within Heliconiini. We focus on five species of Heliconiini, with additional species included where available, which serve as representatives of these groups, to analyse the size and structure of peripheral visual structures, from the eyes, through to the major sensory neuropils, and the major tracts connecting them to the mushroom bodies. These species have overlapping ecological and host-plant preferences (33,43), negating confounding effects of habitat on sensory investment (44,45). We hypothesised that previous evidence for visual specialisation in mushroom body structure and memory stability (30,46) could be reflected in modifications in the visual circuitry peripheral to the mushroom bodies. We show that, despite evidence of visual specialisation in *Heliconius’* mushroom bodies (30), investment in the sensory processing structures and their pattern of downstream projections is largely conserved across Heliconiini.

## Materials and Methods

### i. Eye morphology and imaging

Samples for the analysis of eye structure included a total of 84 adult butterflies (Table S1a), including two representatives of *Heliconius* (21 *Heliconius erato demophoon* and 22 *Heliconius melpomene rosina*) and three from Heliconiini outgroups (19 *Dryas iulia*, 10 *Dryadula phaetusa* and 12 *Agraulis vanillae*). *Dryas iulia* and *Dryadula phaetusa* specimens were collected from stock populations at the Smithsonian Tropical Research Institute insectaries, Gamboa, Panama (May-June 2023). Stocks were established from local populations and maintained in 3×3×2 m mesh cages in ambient conditions and with natural lighting containing pollen resources (*Lantana camara, Palicourea elata* and *Psiguria sp*.) and supplementary artificial feeders containing 20% sugar-water. *Agraulis vanillae* specimens were ordered as pupae from commercial suppliers (The Entomologist; https://butterflypupae.com/), as stocks of this population were not available at the time.

Following eclosion, they were kept in greenhouses in the UK in 2 × 2 × 2 m mesh cages at 26°C, 80% humidity and a 16 h/8 h light/dark cycle. Cages contained supplemental 20% sugar-water as well as *Lantana camara*. All butterflies were cold anesthetised and the head then removed and fixed for 16 hours with agitation in zinc formaldehyde solution (18.4 mM ZnCl2, 135 mM NaCl, 35 mM sucrose, and 1% formaldehyde) **(47)**. The heads were then transferred to Dent’s solution (80% methanol/ 20% DMSO) for 2 hr before being transferred to 100% methanol for -20°C storage.

To quantify variation in eye anatomy, we followed established protocols (48,49). We removed and imaged the eye cuticle, allowing visualisation and quantification of the number and size of the ommatidia present across the eye. Whole heads were imaged using LAS X software and a Leica EZ4 W stereo microscope with a 5 MP camera at 16× magnification. For each individual, the antennae, proboscis and maxillary palps were removed to allow measurement of the interocular distance (IOD), which served as our allometric control (50). Following whole-head imaging, the left eye cuticle was removed for analysis unless it was damaged, in which case the dissection was repeated with the right eye. Once removed, the eye cuticle was placed in 20% NaOH for 16-20 hours to loosen remaining eye tissue behind the cuticular cornea, which was then removed under a 95% ethanol solution (51). Incisions were made along the dorsal-ventral axis and anterior-posterior axis of the cuticle, to allow it to be mounted flat on a microscope slide in Euparal, and left to dry for 24 hours before imaging. Mounted cuticles were imaged using GXCapture software and a Leica M205 C stereo microscope with an integrated camera at 1.6× and 2.0× magnification, depending on cuticle area.

Whole-head and cuticle images were imported into FIJI (52) and scale-corrected. From these images, four morphological variables were recorded: IOD, eye surface area, ommatidia number, and ommatidia size. IOD was taken as the distance between the eyes directly above the proboscis using FIJI’s ‘line tool’. Eye surface area was determined from the cuticle images using the ‘threshold’ and ‘measure’ tool in FIJI (51). Number and size of ommatidia was calculated using the ommatidia detecting algorithm (ODA) python-based pipeline (53).

Using R v4.3.0 (54), linear models were used to analyse inter- and intra-specific differences in eye surface area and ommatidia size across Heliconiini. In all of these models, IOD was included as an allometric control to account for body size differences between species and sexes. Ommatidia diameter, eye surface area and IOD were log10 transformed to meet residual-distribution assumptions of linear models. Linear models were tested sequentially against a null model including only IOD as a control for body size (Table S1B, S1F) to determine best fit. Where differences were found, post-hoc tests with Tukey correction for multiple comparisons were carried out.

### ii. Visual acuity

Using our eye anatomical data, morphological visual acuity was then calculated using ommatidia number following Wright et al. (2023) and based on studies by Land (1997, 1989):

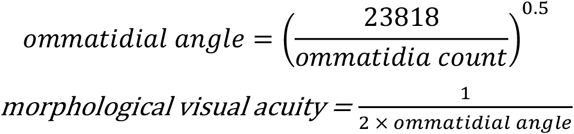

Using R v4.3.0 (54), linear models were used to analyse inter- and intra-specific differences in visual acuity across Heliconiini. Sex was found to be significantly associated with acuity, and was, therefore, accounted for in analyses of interspecific differences. As acuity was directly calculated from ommatidia number, results for visual acuity differences are directly proportional to ommatidia number differences. Visualisation of acuity levels of *Heliconius* and outgroup Heliconiini males and females was carried out in the *AcuityView* v 0.1 package in R (57) on the same visual scene obtained at the forest edge in Gamboa, Panama at four distances (5 m, 10 m, 15 m and 20 m) using a Nikon D3500 (AF-P DX NIKKOR 18-55mm f/3.5-5.6G VR lens). Actual image size was calculated in Fiji (52) using a 1 m pole present in the visual scene.

### iii. Neuropil staining, imaging and segmentation

To examine investment in sensory neuropils, we segmented brain images from 10 *H. melpomene*, 10 *H. erato*, 14 *D. iulia*, 7 *D. phaetusa* and 11 *A. vanillae*. These images were previously obtained from specimens collected from across French Guiana, Panama and Ecuador by Couto et al. (30). Imaging was performed using immunohistochemical staining against a synaptic marker and confocal microscopy, using a standardised protocol. Full details of staining and imaging methods are available in Couto et al. (30) and Montgomery et al. (58). The antennal lobes (AL), medulla (ME), ventral lobula (vLO), the mushroom body calyx, peduncle and lobes, and the rest of the central brain (rCBR) had been previously segmented by Couto et al. (30). For *Heliconius erato* and *Heliconius melpomene*, the additional optic neuropils, the lamina (LA), lobula (LO), lobula plate (LOP), accessory medulla (AME) and the anterior optic tubercule (AOTU), had also been segmented (19).

Here, using Amira 2021.1 (ThermoFisher Scientific, MA, USA), the optic lobe neuropils in *D. iulia, D. phaetusa* and *A. vanillae* were segmented using the ‘labelfield’ module to construct outlines of each structure. For larger structures (LAM, LO and LOP), every fifth image in the stack was manually segmented before interpolating between the images using the ‘interpolation tool’. The interpolated layers were then edited to check for deviations from the staining boundaries and for consistency across the xy, xz and yz views. The volumes of the respective neuropils were extracted using the ‘measure statistics’ module and the 3D models generated using the ‘SurfaceGen’ module. Neuropils were only segmented for one of the brain hemispheres, owing to the consistency in volumes found between each side, and then doubled for statistical analysis (58).

Statistical analyses of investment in the sensory neuropils was carried out using linear models in R v4.3.0 (54) to test for divergence in investment between *Heliconius* and outgroups Heliconiini in scaling and volume of the neuropils compared to the rest of the brain. The volumes of all structures were log10 transformed, and rCBR was used as an allometric control. One *A. vanillae* was removed from the dataset following data visualisation, as it was a clear and consistent outlier in terms of overall size (∼ 3× smaller overall than all other *A. vanillae* in the dataset). Each of the sensory neuropils (LAM, ME, LO, LOP, aME, vLO, AOTU and AL) were analysed separately, testing for an effect of group membership (*Heliconius* vs. all non-*Heliconius* Heliconiini outgroups), as well as the influence of sex differences. Following p-adjustment for multiple testing, sex was found to be insignificant across all structures. Model fit was determined through sequential comparisons to a null-model including only the focal neuropil and rCBR (Table S2a, S2c). The package *emmeans* v1.8.6 (59) was used for post-hoc pairwise comparisons between species, using the estimated marginal means with Tukey correction for multiple comparisons. Allometric scaling relationships between neuropils were analysed using *sma* from the *smatr* v 3.4-8 package in R (60).

### iv. Sensory projection pathways

Finally, to understand how sensory information is relayed to downstream neuropils, particularly the mushroom body, we used injections to mass label projection pathways from the injection site to the terminal sites of the intersected projection neurons, and immunohistochemistry, which were then reconstructed. We made use of samples initially produced for a previous study (30), where tracts from the antennal lobe and optic lobe were stained in two *Heliconius* species (*H. melpomene* and *Heliconius hortense*) and three Heliconiini outgroup species (*D. iulia, A. vanillae* and *Eueides isabella)* using Dextran-conjugated dyes (Fluoro-ruby: 10,000 MW, D1817, Thermo Fisher Scientific; and Alexa fluor 647; 10,000 MW, D22914, Thermo Fisher Scientific) to visualise terminal projection sites within the mushroom body calyx. In brief, these butterflies were immobilised and dye was injected into the antennal lobe, and the dorso-medial optic lobe (targeting the vLO and LO), using a stretched capillary tip and the head was submerged in Ringer solution (30). The dye was left to migrate overnight before the brain was dissected out and fixed. Tracts to the calyx were also stained using retrograde tracing through injection into the mushroom body calyx itself in *H. melpomene* and *D. iulia*. We then mounted the brains in methyl salicylate and imaged them using a laser-scanning microscope (Leica SP8) with a 10× dry objective (NA = 0.4). Here, we re-scanned these samples to provide better contrast and resolution of the sensory pathways upstream of the mushroom body.

To confirm the projection pathways, we used available images of stains of antibodies targeting Glutamate decarboxylase (GAD), acetylated tubulin and Horseradish peroxidase (HRP) staining from in Farnworth et al. (31). Tubulin stains neuronal axons, allowing for visualisation of the neural pathways between neuropils, and GAD labels GABAergic neurons which provide staining of specific cell groups and structural information. Full protocols are available in Farnworth et al. (31), but in brief, *H. melpomene* and *D. iulia* were stained following fixation and sectioning (mounting in Agarose and sectioned using Leica Vibratome VT1000-S), mounted in 80% glycerol on frosted object slides, and imaged using a laser-scanning microscope (Leica SP8) with a 20× dry objective (NA 0.75).

Projection pathways were described and compared between *Heliconius* (*H. melpomene* and *H. hortense*) and outgroup Heliconiini species (*D. iulia, A. vanillae* and *E. isabella*). We focused on the analysis of qualitative differences in the patterns of projections using these stainings. Tract and commissure nomenclature varies between insect studies owing to different brain-body axis and naming systems. Here, the nomenclature used follows Ito et al. (2014), with the addition of the ventral lobula (vLO). Orientation of the brain structures is described based on the body axis. The different staining methods (optic lobe, antennal lobe and calyx injections, and GAD-Tubulin staining) were combined to cross-identify each of the pathways present across Heliconiini species. The clearest representation of each of the pathways from an outgroup and *Heliconius* species is used in the figures (Figure 3, S3, S4) to illustrate the identified tracts and commissures.

## 3. Results

### i. Eye anatomy and visual acuity is largely conserved across Heliconiini

To explore evidence of visual specialisation in *Heliconius*, we first examined whether Heliconiini species differed in ommatidia size. A model including *species, IOD* (as an allometric control) and *eye surface area* showed that ommatidia size varied at both the species level (F4,76 = 24.169, p < 0.001), and in association with surface area (F4,76 = 10.579, p = 0.002). The outgroup Heliconiini species, *A. vanillae*, had larger ommatidia than all other species, with *D. phaetusa* also having larger ommatidia than *H. erato* (Table S1Ciii). In a model including a group level factor rather than species, differences were found between *Heliconius* and outgroup Heliconiini (F1,79 = 4.638, p = 0.034), but with smaller diameters on average in *Heliconius*. Across both the species level (F4,76 = 14.980, p < 0.001) and group level (F1,79 = 18.051, p < 0.001) analyses, males had significantly higher average ommatidia diameter than females, apart from in *H. melpomene*, where intraspecific sexual dimorphism in ommatidia size was not detected (Figure 1B, Table S1Cii).

**Figure 1.**
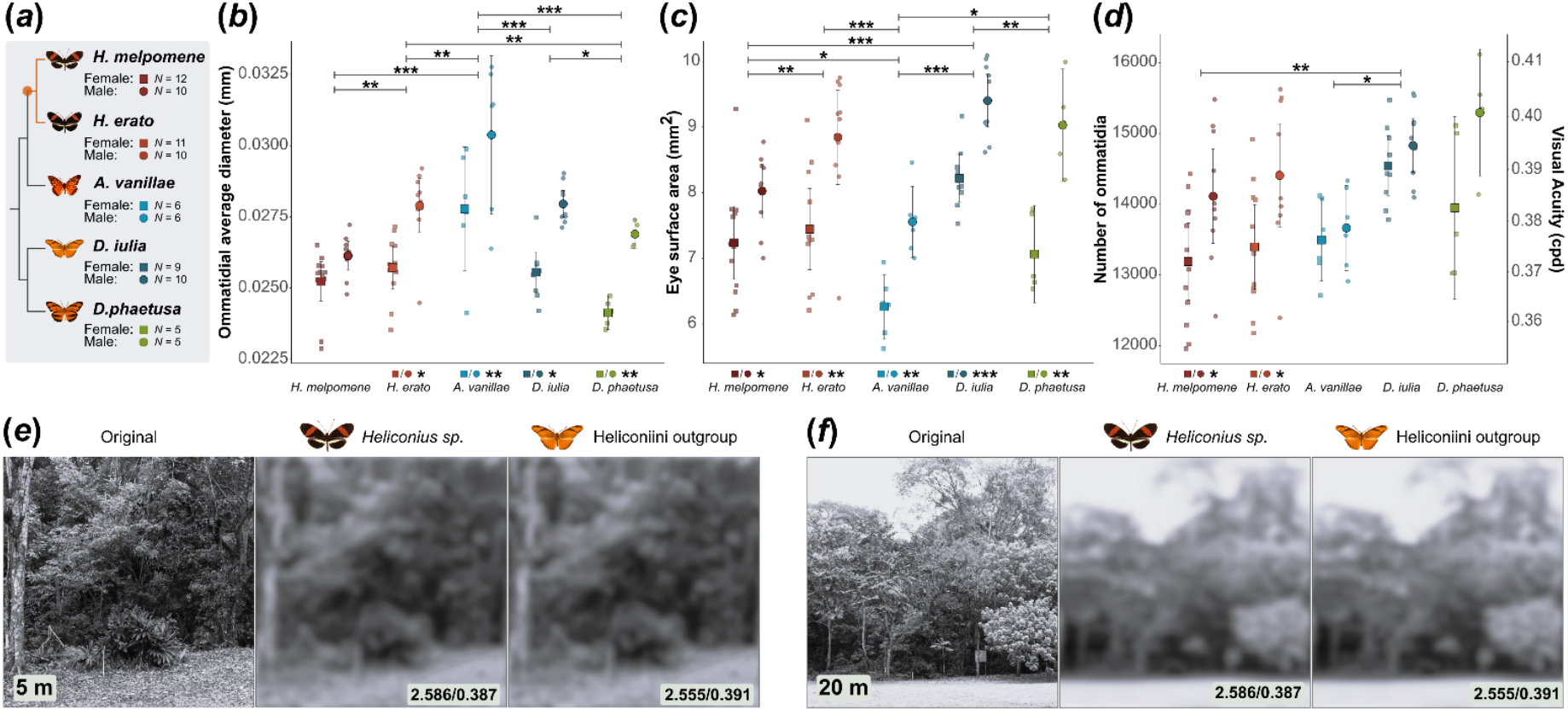
Eye morphology and visual acuity across *Heliconius* and outgroup Heliconiini. **(a)** Heliconiini phylogeny with pollen feeding innovation indicated in orange. **(b)** Average ommatidia diameter differences between Heliconiini species male (circle) and females (square). Significance bars are species differences in surface area (See Table S1c for variation with and without accounting for surface area). **(c)** Surface area (mm^2^) differences between Heliconiini species and males (circle) and females (square). **(d)** Acuity is directly calculated from ommatidia number which are plotted on the right axis and morphological visual acuity on the left axis. Squares represent female acuity means and circles female acuity means with lines showing the confidence intervals. **(e/f)** Mean morphological visual acuity in male Heliconiini demonstrating ability to distinguish main canopy features at distances of 5 m and 20 m (See Figure S1 for additional visualisations). Visualisations of the scenery were generated with the AcuityView package in R. Values show morphological estimates of the minimum resolvable angle in degree/ acuity in cycles-per-degree. Significant differences between species are indicated by significance bars. Significant sex differences are indicated above each species label. Group differences between *Heliconius* and Heliconiini outgroup are indicated in the bottom right of each plot. (*p < 0.05, **p < 0.01, ***p < 0.001).

Next, we analysed variation in eye surface area overall, and found there to be considerable interspecific differences (F4,77 = 16.264, p < 0.001; Table S1Div). However, these were consistent within groups, and as such group level differences between *Heliconius* and outgroup Heliconiini species were not significant (F1,80 = 0.094, p = 0.761). Despite this, we still identified pronounced variation found between most species (Table S1Fv) including within the *Heliconius* genus, and among outgroup Heliconiini. Surface area also varied significantly with sex at both the species level (F4,77 = 81.714, p < 0.001) and group level (F1,80 = 46.652, p < 0.001), with males consistently possessing greater eye surface area than females (Figure 1C). Finally, we analysed differences in ommatidia count between species, groups and sexes. Species differences (F4,77 = 5.186, p < 0.001), though minimal (see Table S1Bvi for post-hoc comparisons), as well as group differences (F1,80 = 4.618, p = 0.035) were found in ommatidia count. Overall, the outgroup Heliconiini group was found to have higher ommatidia numbers than *Heliconius* (Figure 1D). Between-species post-hoc analyses revealed that there were only differences between *A. vanillae* and *D. iulia*, and *D. iulia* and *H. melpomene*. Sex was also found to significantly vary with ommatidia count, at both the group (F1,80 = 18.998, p < 0.001) and species level (F1,77 = 21.286, p < 0.001), with males having a higher number of ommatidia. However, when looking at intraspecific sex differences, these were isolated to *H. erato* and *H. melpomene* (Figure 1D). Despite this, an interaction between sex and species, or sex and group, was not found to be significant (Table S1Biv, S1Fix). In summary, based on our morphological measures of ommatidia size and number, *Heliconius* do not have more structurally expanded eyes than outgroup Heliconiini.

To explore how these anatomical traits reflect visual acuity, we used ommatidia count to estimate visual acuity capacity across the species (55,56), as well as the *AcuityView* package to visualise their perception of visual canopy scenes from different distances. As expected given the above results, we found evidence of interspecific (F4,77 = 5.230, p < 0.001) and intergroup variation (F1,80 = 4.510, p = 0.037), as well as sexual dimorphism in visual acuity (Figure 1D, Table S1F, S1B) that closely mirrored the ommatidia count results. Again, no significant interaction between sex and species was found (Table S1Bi, S1Fi, S1Fii) and intraspecific sex differences in acuity were isolated to *Heliconius* species (Table S1Bii). Visualisations of variation in visual acuity revealed that both groups can determine major canopy features in their environment from a range of distances between 5 m and 20 m

(Figure 1E/F, S1), but no difference between *Heliconius* and the Heliconiini outgroups were evident.

### ii. Heliconius *do not invest more in visual processing centres of the optic lobes*

Measurements of eye anatomy suggest *Heliconius* capture visual information similarly to Heliconiini outgroups. We therefore assessed whether *Heliconius* have increased investment in the processing of visual cues through increases in the size or one or more of the six visual neuropils (Figure 2). Sex did not have an explanatory relationship with sensory neuropil volumes (Table S2Ai) and was, therefore, excluded as a variable in further analyses. The volume of the antennal lobe, the primary olfactory neuropil, was conserved across species (F4,45= 1.396, p = 0.251; Figure 2G). In contrast, interspecific variation was found in seven of the eight neuropils (Table S2Aii). In the majority of these cases, interspecific differences were a result of increased investment in the non-*Heliconius* Heliconiini outgroups rather than *Heliconius* species. The lamina (F1,41= 44.717, p < 0.001), lobula (F1,48= 21.460, p < 0.001) and AOTU (F1,48= 30.338, p < 0.001) were all significantly larger in the outgroups compared to *Heliconius* (Table S2c, Figure S2). In contrast, at the group level the ventral lobula was larger in *Heliconius* than the outgroups (F1,48= 19.724, p < 0.001; Figure S2), but was consistent between *Heliconius* and the outgroup species *A. vanillae* when examining species-specific differences, suggesting a degree of overlap between the groups (Table S2b). Comparisons of allometric scaling relationships further confirmed these results, highlighting non-allometric grade-shifts between species, with the exception of a major axis shift in the antennal lobe (Figure 2G, Table S3Ai). Where grade-shifts were present, most reflected either an increased or conserved investment in the outgroup species relative to *Heliconius* species (Table S3Aii; Figure 2D-F, Figure S2). In fact, across most of the neuropils, *D. phaetusa* and *D. iulia* investment was significantly higher than other species, with neuropil volume largely conserved between these two species (Table S3Aii; see supplementary text for further details), which is consistent with their phylogenetic relatedness. Ultimately, these results show there is no consistent, significant pattern of enlargement in the sensory neuropils in *Heliconius* that is coincident with the expansion and visual specialisation of the mushroom body (30)

**Figure 2.**
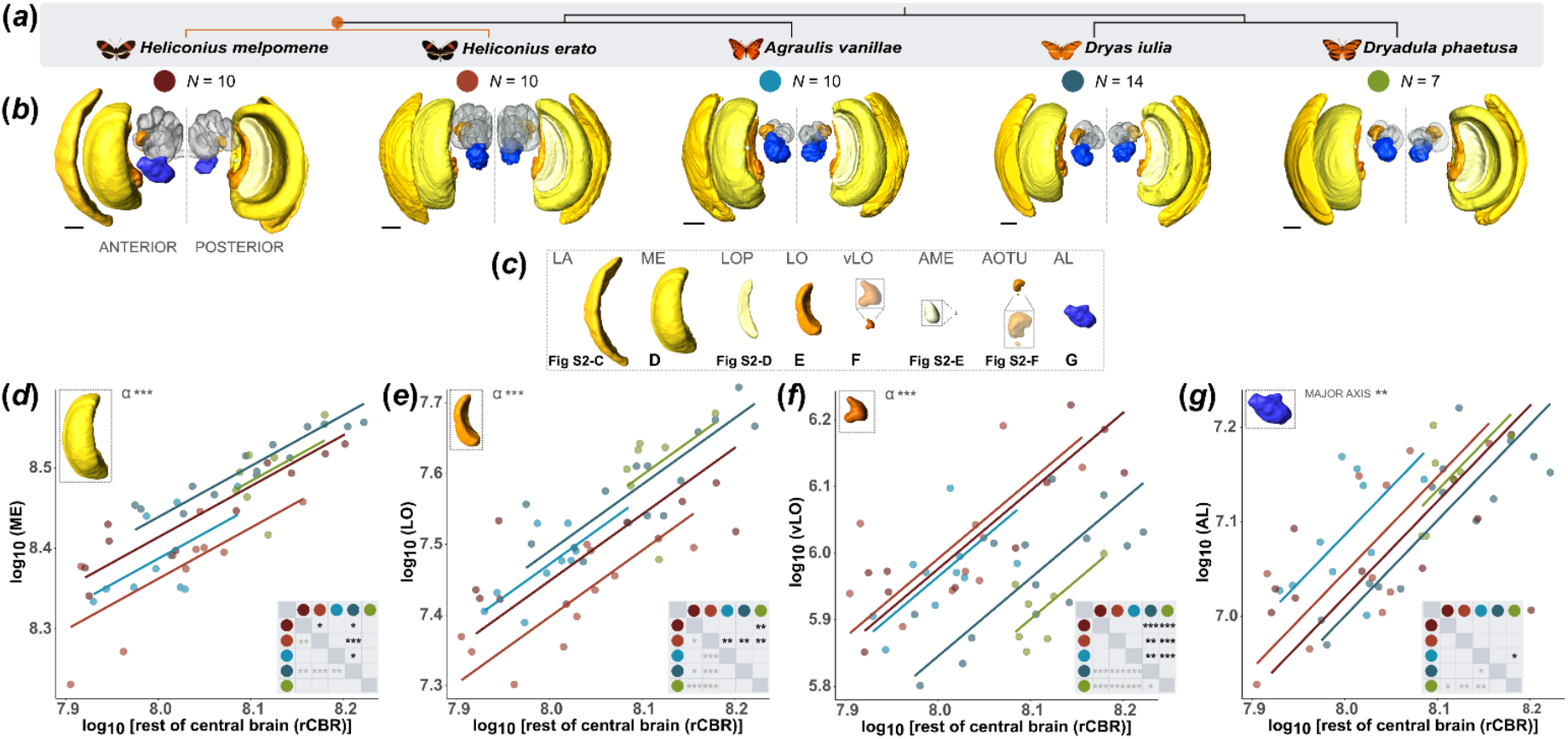
Sensory neuropil investment across Heliconiini. (**a**) Heliconiini phylogeny with evolution of pollen feeding in *Heliconius* indicated in orange. (**b**) Anterior (left) and posterior (right) view of 3D surface reconstructed sensory neuropils. Neuropil in yellow-orange: visual neuropil, grey: mushroom bodies, blue: antennal lobes. (**c**) Surface reconstructions of the sensory neuropils moving from the periphery (left) to central brain (right). (**d-g**) Scaling relationships between four of the sensory neuropils (AL, ME, LO and vLO) and the rest of the central brain (rCBR). Red: *Heliconius* species, blue/green: non-pollen feeding Heliconiini. Each plot contains a matrix of pairwise species differences in the lower right corner. Level of significance indicated before (light grey) and after p-adjustment (black) (*p < 0.05, **p < 0.01, ***p < 0.001). For analyses of the other additional neuropils and group differences see Figure S2. Scale bars are 200 μm.

### iii. Sensory projection pathways are qualitatively conserved across Heliconiini

In light of the lack of increased investment in visual acuity or visual processing in the primary sensory neuropils in *Heliconius*, we aimed to further understand whether increased visual reliance in the mushroom bodies and visual performance is the result of differences in supply of visual and olfactory information to the mushroom bodies. Generally, our labelling of the projection neurons from the optic lobe were consistent with projection neurons identified in other species (6,40,62). We saw major inputs to neuropils of the central brain, the mushroom bodies, and to the contralateral optic lobe (Figure 3, Figure S3). Here, we focused on projections to the mushroom bodies, which are necessary for sensory integration, and major associated neuropils such as the AOTU, which has important functions in processing skylight cues and encoding of visual components for navigation through the anterior visual pathway (63,64). Additional anatomical descriptions can be found in the supplementary material (Supplementary Results S2).

**Figure 3.**
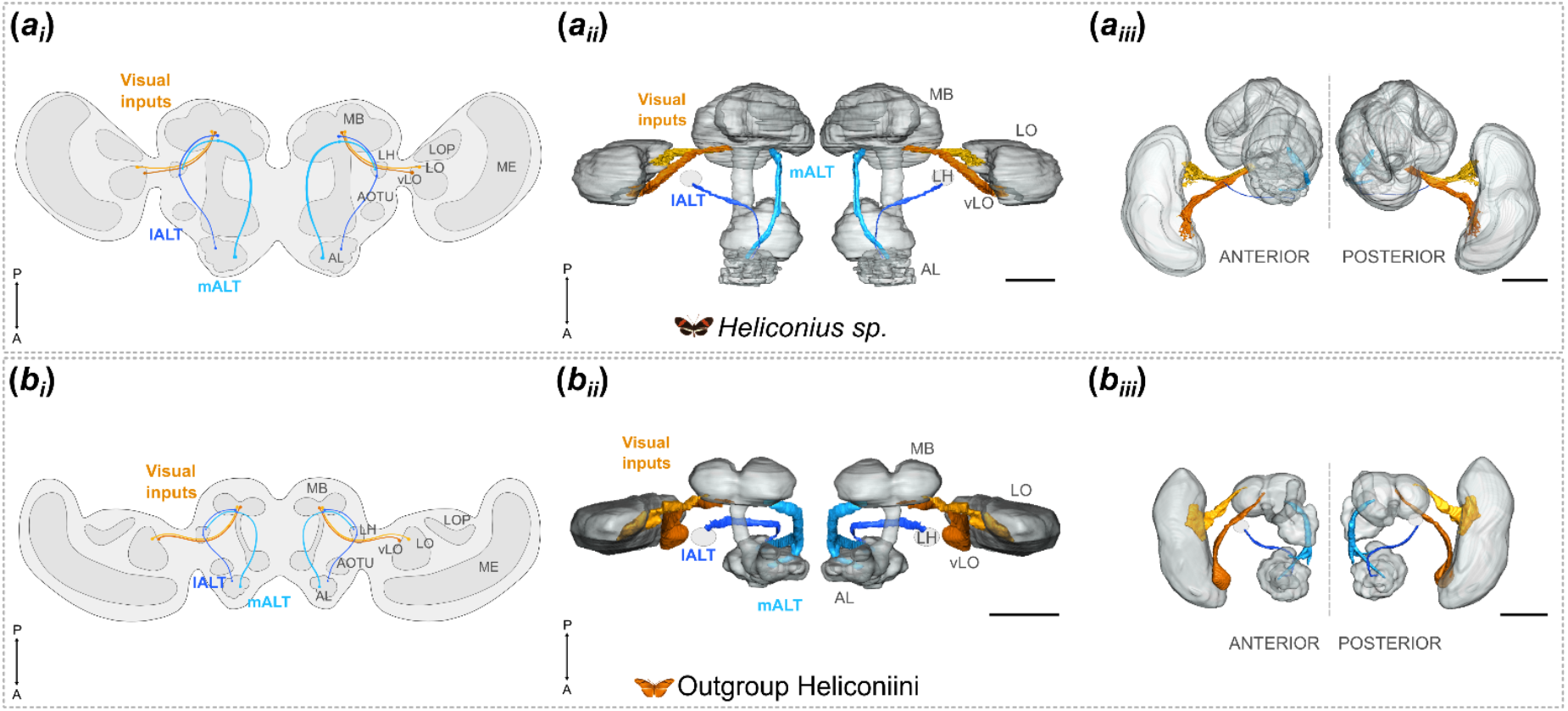
Major visual and olfactory projection pathways in Heliconiini. **(a)** Projection pathways to the mushroom body calyces in *Heliconius*. **(b)** Projection pathways to the mushroom body calyces in outgroup Heliconiini species. **a**_**i**_**/b**_**i**_: Schematic of the two visual projections to the mushroom body calyces originating in the ventral lobula (vLO, orange) and lobula (LO, yellow), and two olfactory projections the medial antennal lobe tract (mALT, light blue) and lateral antennal lobe tract (lALT, dark blue) from the antennal lobe (AL) to the calyces (CA) in *Heliconius* (A_1_) and outgroup Heliconiini (B_1_). **a**_**ii**_**/b**_**ii**_: 3D segmentations of projection pathways to the mushroom bodies and associated visual and olfactory neuropils based on calyx injection in *H. melpomene* (A_2_) and *D. iulia* (B_2_) from a dorsal-ventral and anterior-posterior perspective. The lateral horn (LH) could not be reliably segmented and is therefore represented by a dashed circle. The lALT projects first to the LH and then to the calyx. **a**_**iii**_**/b**_**iii**_: Anterior-posterior perspective of the segmented projection pathways. See figures S3 and S4 for full overview of all projection pathways and pathway images. Scale bars are 200 μm.

In both *Heliconius* and outgroup Heliconiini species, the major input to the mushroom body calyx from the optic lobes was via a thick tract connecting the optic lobe to the ipsilateral calyx (Figure 3; Figure S3). A similar tract has been identified in other Lepidoptera and insect species, but with variation in where the tract originates and pattern of projection. In Heliconiini, calyx injections in *D. iulia* and *H. melpomene* revealed this tract to be composed of projections from the lobula and the ventral lobula that converge in the central brain before reaching and innervating the base of the calyx. Fibres from across the lobula converge medially to form the more dorsal lobula tract from the lobula to the calyx (Figure 3, S3). The ventral lobula tract projects dorsally from the ventral lobula into the central brain where it converges with the lobula tract. In addition to the visual projection to the calyx, a prominent thick projection, the anterior optic tract (AOT), connects the lobula with the AOTU, and from the lower AOTU subunit some neurons project further to the lateral complex (Figure S3). This tract was observed in both *Heliconius* and the outgroup Heliconiini species and is located anteriorly to the tracts that connect the optic lobe and calyx (Figure S3). The tubercle to tubercle tract (TUTUT) connected the ipsilateral and contralateral AOTU across the hemispheres (Figure S3).

We also considered whether the disproportionate expansion of the visual calyx may reflect a reduction of projection neurons from the antennal lobe. In both *Heliconius* and their outgroup genera, we identified a pattern of four major olfactory projections from the antennal lobes, as observed in other insects (6,41,65). Injections into the calyx reveal that two of the four antennal lobe tracts project to the calyx (Figure 3). The medial antennal lobe tract (mALT), projects from the antennal lobe posteriorly and dorsally where it sends branches directly to the calyx, before projecting laterally to the lateral horn (Figure 3). The lateral antennal lobe tract (lALT), projects first laterally from the antennal lobe up to the lateral horn before a small number of fibres innervate the calyx. This tract was present in both antennal lobe and calyx injections, with the calyx injections revealing that only a small number of neurons in the lALT project further from the lateral horn to the calyx (Figure 3), with the majority terminating in the lateral horn, revealed through antennal lobe injections. The mediolateral antennal lobe tract (mlALT; 58) and transverse antennal lobe tract (tALT; 41), both project from the antennal lobe to the lateral horn (Figure S4). The orientation of these two tracts varied slightly between *Heliconius* and the outgroup Heliconiini, owing to the expansion of the mushroom bodies in *Heliconius*. Generally, however, qualitative comparisons of both sensory and olfactory projection pathways downstream of the mushroom body reveal a high degree of conservation across Heliconiini in the supply channels leading to the mushroom bodies.

## 4. Discussion

Increased mushroom body size and Kenyon cell number in the genus *Heliconius* is strongly linked to the evolution of a cognitively demanding foraging behaviour associated with pollen feeding (30). Furthermore, expansion of the mushroom bodies largely reflects an increase in the volume of the visual mushroom body calyx (30) and lobes (31), which, along with evidence of improved long-term visual memory (46), suggests a strong degree of visual specialisation in learning and memory circuits. However, our data illustrating conservation of the sensory structures and pathways across multiple Heliconiini support an isolated change in the *Heliconius* mushroom bodies in facilitating this behavioural shift, while also providing a previously uncharacterised map of key projection pathways within the Heliconiini brain. Our analyses of eye anatomy, sensory neuropil volume and pattern of sensory tracts reveal that the evolutionary modification of the neural circuitry in *Heliconius* is seemingly highly localised to the mushroom bodies, with the sensory structures peripheral to the mushroom bodies found to be highly conserved between *Heliconius* and the outgroup Heliconiini. Below, we discuss each of the components analysed starting from the periphery and ending with the projection pathways innervating the mushroom bodies.

Despite *Heliconius*’ capacity for spatial learning (24,25), improved visual memory (27), and expanded visual calyx (30), we found no evidence for *Heliconius*-specific increases in visual acuity. Indeed, the Heliconiini outgroups had higher acuity than the *Heliconius* genus, which indicates that spatial foraging does not require increased resolution of visual scenes, but rather differences in how these senses are processed in integration centres such as the mushroom bodies (30), and potentially the central complex (66). It is possible that the evolution of higher acuity in the outgroups may be explained by factors such as flight dynamics or differences in motion detection, which also correspond with ommatidia number and size across the eye (55,67,68). Despite this, visualisation of *Heliconius*’ ability to perceive details of visual scenes suggests that landscapes could be resolved in sufficient degree for their use in guiding spatial learning (24). Our data build on previous research on acuity and eye morphology within *Heliconius* (37,48,51,69,70), but importantly, provide new data for outgroup Heliconiini.

We note that our acuity estimates for the eye are based on structural differences in the eye cuticle, which does not fully account for the downstream processing in the eye the way behavioural assessments of acuity would (55,71). Indeed, behavioural assays produce increased estimates of acuity over morphological measures, although they also mirror intergroup patterns seen in morphological tests (51), suggesting they remain a useful indicator of relative performance. Additionally, if processing differences were present between the *Heliconius* and outgroup species, this would likely be represented in the volumes of sensory neuropils, in particular, the lamina and medulla, which receive the main direct photoreceptor input from the eye (72). Although neuropil investment differences were found between species, there was no consistent pattern of variation between species that would explain the behavioural differences seen in *Heliconius*. In fact, contrary to prior expectations, greater investment was consistently seen in the outgroup Heliconiini across the sensory neuropils. The exception to this was the ventral lobula where, overall, *Heliconius* had greater investment. The ventral lobula is a visual neuropil closely linked to the mushroom bodies (39,40), and, as confirmed by the pathway analysis, one of the major mushroom body inward projection tracts originates from this area (30). The significant grade shift found in *Heliconius* specifically would, therefore, align with the expansion of the visual calyx. However, when differences were assessed at a species level, *Heliconius* had consistent investment with the outgroup Heliconiini species *A. vanillae*. Furthermore, a previous study of a wider range of *Heliconius* and outgroup Heliconiini species also found that expansion of the ventral lobula was not unique to *Heliconius* (30). As a result, we suggest that there is limited evidence that greater investment in the visual neuropils facilitates the systematic foraging seen in *Heliconius*.

We also found no obvious differences in the sensory projections to the mushroom bodies and other integrative neuropils. Both the visual and olfactory projections found were both highly conserved within the *Heliconius* and non-*Heliconius* Heliconiini, as well as consistent with sensory pathways observed in other Lepidoptera (40,41,73) and even other insect orders (74). Although the diameter of the projection pathways were not quantified, we did not observe any obvious qualitative variation in the parameters of the tracts between species. This suggests a potential difference in the degree of Kenyon cell sampling of the visual projection neurons between the *Heliconius* and outgroup Heliconiini, given the increase in Kenyon cell number but no obvious increase in visual projection pathway density (30,75). Segregation of the visual and olfactory inputs to the calyces has been demonstrated across Heliconiini genera (30), as well as in other species within and outside of Lepidoptera (40,76,77). Indeed, visual input to the mushroom bodies in Heliconiini is similar to that observed in the swallowtail, *Papilio xuthus*, with regards to the projection neurons originating in both the ventral lobula and lobula (40). Projection neurons from the lobula to the calyx are commonly seen in visually orientated insect species, with species in Hymenoptera exhibiting both lobula and medulla input (62,78–80). In Heliconiini, however, there were no clear indications of direct input from the medulla to the calyx or AOTU via the AOT from the injections into either the calyx or the optic lobes. This does not confirm their absence, but the lack of staining observed in the medulla compared to the strong presence of the lobula complex suggests there are, perhaps, different integration needs in Heliconiini in comparison to other insect and Lepidoptera species. Olfactory input to the calyces in Heliconiini was also consistent with that seen in other Lepidoptera, with two tracts from the antennal lobes (mALT and lALT) projecting to the mushroom bodies (40,41,65) and the patterns and apparent size of olfactory projections conserved between the *Heliconius* and outgroup Heliconiini species. Although analysis of the tracts was largely observational through 3D reconstruction and image analysis, and the mass staining methodology may be obscuring finer, detailed differences, the lack of obvious differences suggests any changes between *Heliconius* and the outgroup Heliconiini are minor in comparison to the large changes seen in the mushroom bodies.

Although not the primary focus of the current study, our results on eye structure suggest divergent patterns of sexual dimorphism in specific traits. Sexual dimorphism was only detected in the number and size of ommatidia, and not in investment in downstream neuropils, suggesting a primary importance in acuity and light capture. We also found consistent sexual dimorphism in ommatidia number, and therefore visual acuity, between *Heliconius* and their outgroups. Sexual dimorphism was only significant in *Heliconius* species, as previously observed (37,51), but was not evident in the three outgroup species. In contrast, ommatidia size showed interspecific differences in levels of sexual dimorphism, but not consistently between *Heliconius* and outgroups. As such, sexual dimorphism in visual acuity in *Heliconius* may be the result of *Heliconius* males actively seeking out mates (51).

Mating behaviour in at least one outgroup species, *Dryas iulia*, has been suggested to involve less complex recognition of conspecifics than in *Heliconius* (81), supporting a potential link between mating behaviour and increased male visual acuity in *Heliconius*. Notably, sexual dimorphism has previously been described in the retinal mosaic of *Heliconius* eyes, reflecting sex differences in the combinations of opsins expressed within an ommatidia among species in the Erato and Sara/Sapho groups (36,82,83). Although the outgroup genera have not been studied in depth, *Euiedes isabella* has a sexually monomorphic retinal mosaic (83). As such, our data on ommatidia number adds to evidence that the *Heliconius* eye may be adapted to meet the demands of sex-specific behaviours.

Changes in ommatidia size may instead be indicative of selection towards greater photon capture in male Heliconiini (69), but not in a manner that distinguishes *Heliconius* from outgroups.

In summary, contrary to prior expectations, we found no increased investment in the sensory structures peripheral to the mushroom bodies in *Heliconius* that would facilitate their systematic foraging and increased visual-memory stability. Surprisingly, whilst there are still open questions with regards to the morphology and associated cells of the sensory projection neurons, *Heliconius* showed conserved, or even lower, investment in the visual domain compared to the outgroup Heliconiini. We therefore infer that the expansion of the mushroom bodies in *Heliconius*, and the associated cognitive enhancements, are the result of circuit restructuring, particularly in regard to the synaptic fields within the mushroom bodies (1,2). This is supported by previous evidence of increased synaptic plasticity in the mushroom bodies of *Heliconius* over the outgroup Heliconiini species (27). The restructuring of these learning and memory circuits may be facilitating a shift in the weighting of Kenyon cell input sites, allowing for evolutionary changes in specific functional domains. Additionally, the conservation of the sensory structures suggests that sufficient sensory processing had already evolved prior to the split of *Heliconius* from Heliconiini, and instead highlights the importance of the evolution of visual learning and long-term memory as primary adaptations underpinning behavioural innovation in *Heliconius* foraging behaviour. Heliconiini, therefore, present an interesting case of genus-specific mosaic brain expansion allowing exploitation of a new niche, furthering our understanding of cognitive evolution.

## Acknowledgements

We thank the Ministerio del Ambiente, Panama, and research and facilities staff at the Smithsonian Tropical Research Institute in Panama, in particular Rémi Mauxion, Oscar Paneso, Cruz Batista, for facilitating our work. We thank John Currea for the assistance with running the Ommatidia Detecting Algorithm and helpful discussions about the research.

Additionally, we gratefully acknowledge the Wolfson Bioimaging Facility for imaging and analysis support. This work was supported by a NERC Independent Research Fellowship (NE/N014936/1) and an ERC Starter Grant (758508) to SHM and the Smithsonian Tropical Research Institute.

## Bibliography

1. Farnworth MS, Montgomery SH. Evolution of neural circuitry and cognition. Biology Letters. 2024 May 15;20(5):20230576.

2. Roberts RJV, Pop S, Prieto-Godino LL. Evolution of central neural circuits: state of the art and perspectives. Nat Rev Neurosci. 2022 Dec;23(12):725–43.

3. Dell’Aglio DD, McMillan WO, Montgomery SH. Shifting balances in the weighting of sensory modalities are predicted by divergence in brain morphology in incipient species of Heliconius butterflies. Animal Behaviour. 2022 Mar 1;185:83–90.

4. Montgomery SH, Ott SR. Brain composition in Godyris zavaleta, a diurnal butterfly, Reflects an increased reliance on olfactory information. Journal of Comparative Neurology. 2015;523(6):869–91.

5. Stöckl A, Heinze S, Charalabidis A, el Jundi B, Warrant E, Kelber A. Differential investment in visual and olfactory brain areas reflects behavioural choices in hawk moths. Sci Rep. 2016 May 17;6(1):26041.

6. Immonen EV, Dacke M, Heinze S, el Jundi B. Anatomical organization of the brain of a diurnal and a nocturnal dung beetle. Journal of Comparative Neurology. 2017;525(8):1879–908.

7. Carlson BA, Hasan SM, Hollmann M, Miller DB, Harmon LJ, Arnegard ME. Brain Evolution Triggers Increased Diversification of Electric Fishes. Science. 2011 Apr 29;332(6029):583–6.

8. Ehmer B, Gronenberg W. Mushroom body volumes and visual interneurons in ants: Comparison between sexes and castes. Journal of Comparative Neurology. 2004;469(2):198–213.

9. Hong RL, Riebesell M, Bumbarger DJ, Cook SJ, Carstensen HR, Sarpolaki T, et al. Evolution of neuronal anatomy and circuitry in two highly divergent nematode species. Sengupta P, Marder E, editors. eLife. 2019 Sep 17;8:e47155.

10. Seeholzer LF, Seppo M, Stern DL, Ruta V. Evolution of a central neural circuit underlies Drosophila mate preferences. Nature. 2018 Jul;559(7715):564–9.

11. Gilbert LE. Pollen Feeding and Reproductive Biology of Heliconius Butterflies. Proc Natl Acad Sci USA. 1972 Jun;69(6):1403–7.

12. Hikl AL, Krenn HW. Pollen Processing Behavior of Heliconius Butterflies: A Derived Grooming Behavior. Journal of Insect Science. 2011 Aug 9;11(1):99.

13. Jiggins CD. The passion: niche differentiation, coexistence, and coevolution. In: Jiggins CD, Lamas G, editors. The Ecology and Evolution of Heliconius Butterflies [Internet]. Oxford University Press; 2016 [cited 2025 May 1]. p. 0. Available from: 10.1093/acprof:oso/9780199566570.003.0003

14. Cardoso MZ. Patterns of pollen collection and flower visitation by Heliconius butterflies in southeastern Mexico. Journal of Tropical Ecology. 2001 Sep;17(5):763–8.

15. Dunlap-Pianka HL. Ovarian dynamics in Heliconius butterflies: Correlations among daily oviposition rates, egg weights, and quantitative aspects of oögenesis. Journal of Insect Physiology. 1979 Jan 1;25(9):741–9.

16. O’Brien DM, Boggs CL, Fogel ML. Pollen feeding in the butterfly Heliconius charitonia: isotopic evidence for essential amino acid transfer from pollen to eggs. Proceedings of the Royal Society of London Series B: Biological Sciences. 2003 Dec 22;270(1533):2631–6.

17. Turner JRG. Experiments on the Demography of Tropical Butterflies. II. Longevity and Home-Range Behaviour in Heliconius erato. Biotropica. 1971;3(1):21–31.

18. Pinheiro de Castro EC, McPherson J, Jullian G, Mattila ALK, Bak S, Montgomery SH, et al. Pollen-feeding delays reproductive senescence and maintains toxicity of Heliconius erato. Peer Community Journal [Internet]. 2025 [cited 2025 May 27];5. Available from: https://peercommunityjournal.org/articles/10.24072/pcjournal.546/

19. Hebberecht L, Melo-Flórez L, Young FJ, McMillan WO, Montgomery SH. The evolution of adult pollen feeding did not alter postembryonic growth in Heliconius butterflies. Ecology and Evolution. 2022;12(6):e8999.

20. Young FJ, Montgomery SH. Pollen feeding in Heliconius butterflies: the singular evolution of an adaptive suite. Proceedings of the Royal Society B: Biological Sciences. 2020 Nov 11;287(1938):20201304.

21. Young FJ, Montgomery SH. Heliconiini butterflies as a case study in evolutionary cognitive ecology: behavioural innovation and mushroom body expansion. Behav Ecol Sociobiol. 2023 Nov 28;77(12):131.

22. Lihoreau M, Raine NE, Reynolds AM, Stelzer RJ, Lim KS, Smith AD, et al. Unravelling the mechanisms of trapline foraging in bees. Commun Integr Biol. 2013 Jan 1;6(1):e22701.

23. Menzel R, Greggers U. The memory structure of navigation in honeybees. J Comp Physiol A. 2015 Jun 1;201(6):547–61.

24. Moura PA, Cardoso MZ, Montgomery SH. Heliconius butterflies use wide-field landscape features, but not individual local landmarks, during spatial learning. Royal Society Open Science. 2024 Nov 6;11(11):241097.

25. Moura PA, Young FJ, Monllor M, Cardoso MZ, Montgomery SH. Long-term spatial memory across large spatial scales in Heliconius butterflies. Curr Biol. 2023 Aug 7;33(15):R797–8.

26. Moura PA, Corso G, Montgomery SH, Cardoso MZ. True site fidelity in pollen-feeding butterflies. Functional Ecology. 2022;36(3):572–82.

27. Young FJ, Alcalde A, Melo-Flórez L, Couto A, Foley J, Monllor M, et al. Enhanced long-term memory and increased mushroom body plasticity in Heliconius butterflies. iScience. 2024 Jan 18;108949.

28. Hodge EA, Alcalde Anton A, Bestea L, Hernández G, Margareth Aguilar J, Farnworth MS, et al. Modality-specific long-term memory enhancement in Heliconius butterflies. Philosophical Transactions of the Royal Society B: Biological Sciences. 2025 Jun 26;380(1929):20240119.

29. Zars T. Behavioral functions of the insect mushroom bodies. Current Opinion in Neurobiology. 2000 Dec 1;10(6):790–5.

30. Couto A, Young FJ, Atzeni D, Marty S, Melo-Flórez L, Hebberecht L, et al. Rapid expansion and visual specialisation of learning and memory centres in the brains of Heliconiini butterflies. Nat Commun. 2023 Jul 7;14(1):4024.

31. Farnworth MS, Loupasaki T, Couto A, Montgomery SH. Mosaic evolution of a learning and memory circuit in Heliconiini butterflies. Curr Biol. 2024 Nov 18;34(22):5252-5262.e5.

32. Cicconardi F, Milanetti E, Pinheiro de Castro EC, Mazo-Vargas A, Van Belleghem SM, Ruggieri AA, et al. Evolutionary dynamics of genome size and content during the adaptive radiation of Heliconiini butterflies. Nat Commun. 2023 Sep 12;14(1):5620.

33. Brown KS. The Biology of Heliconius and Related Genera. Annu Rev Entomol. 1981 Jan;26(1):427–57.

34. Finkbeiner SD, Briscoe AD. True UV color vision in a female butterfly with two UV opsins. Journal of Experimental Biology. 2021 Sep 29;224(18):jeb242802.

35. McCulloch KJ, Macias-Muñoz A, Mortazavi A, Briscoe AD. Multiple Mechanisms of Photoreceptor Spectral Tuning in Heliconius Butterflies. Molecular Biology and Evolution. 2022 Apr 1;39(4):msac067.

36. McCulloch KJ, Osorio D, Briscoe AD. Sexual dimorphism in the compound eye of Heliconius erato: a nymphalid butterfly with at least five spectral classes of photoreceptor. Journal of Experimental Biology. 2016 Aug 1;219(15):2377–87.

37. Seymoure BM, Mcmillan WO, Rutowski R. Peripheral eye dimensions in Longwing (Heliconius) butterflies vary with body size and sex but not light environment nor mimicry ring. The Journal of Research on the Lepidoptera. 2015;48:83–92.

38. Couto A, Wainwright JB, Morris BJ, Montgomery SH. Linking ecological specialisation to adaptations in butterfly brains and sensory systems. Current Opinion in Insect Science. 2020 Dec 1;42:55–60.

39. Heinze S, Reppert SM. Anatomical basis of sun compass navigation I: The general layout of the monarch butterfly brain. Journal of Comparative Neurology. 2012;520(8):1599–628.

40. Kinoshita M, Shimohigasshi M, Tominaga Y, Arikawa K, Homberg U. Topographically distinct visual and olfactory inputs to the mushroom body in the Swallowtail butterfly, Papilio xuthus. Journal of Comparative Neurology. 2015;523(1):162–82.

41. Ian E, Berg A, Lillevoll SC, Berg BG. Antennal-lobe tracts in the noctuid moth, Heliothis virescens: new anatomical findings. Cell Tissue Res. 2016 Oct 1;366(1):23–35.

42. Kanzaki R, Soo K, Seki Y, Wada S. Projections to Higher Olfactory Centers from Subdivisions of the Antennal Lobe Macroglomerular Complex of the Male Silkmoth. Chemical Senses. 2003 Feb 1;28(2):113–30.

43. Merrill RM, Naisbit RE, Mallet J, Jiggins CD. Ecological and genetic factors influencing the transition between host-use strategies in sympatric Heliconius butterflies. Journal of Evolutionary Biology. 2013 Sep 1;26(9):1959–67.

44. Montgomery SH, Rossi M, McMillan WO, Merrill RM. Neural divergence and hybrid disruption between ecologically isolated Heliconius butterflies. Proceedings of the National Academy of Sciences. 2021 Feb 9;118(6):e2015102118.

45. Montgomery SH, Merrill RM. Divergence in brain composition during the early stages of ecological specialization in Heliconius butterflies. Journal of Evolutionary Biology. 2017;30(3):571–82.

46. Hodge E, Anton AA, Bestea L, Hernández G, Aguilar JM, Farnworth MS, et al. Modality specific memory enhancement in Heliconius butterflies [Internet]. bioRxiv; 2024 [cited 2025 Feb 10]. p. 2024.09.14.612954. Available from: https://www.biorxiv.org/content/10.1101/2024.09.14.612954v1

47. Ott SR. Confocal microscopy in large insect brains: Zinc–formaldehyde fixation improves synapsin immunostaining and preservation of morphology in whole-mounts. Journal of Neuroscience Methods. 2008 Jul 30;172(2):220–30.

48. Wright DS, Rodriguez-Fuentes J, Ammer L, Darragh K, Kuo CY, McMillan WO, et al. Selection drives divergence of eye morphology in sympatric Heliconius butterflies. Evolution. 2024 May 13;qpae073.

49. Wainwright JB, Loupasaki T, Ramírez F, Williams ILP, England SJ, Barker A, et al. Mutualisms within light microhabitats drive sensory convergence in a mimetic butterfly community [Internet]. bioRxiv; 2024 [cited 2025 Jul 1]. p. 2024.08.16.607924. Available from: https://www.biorxiv.org/content/10.1101/2024.08.16.607924v2

50. Wainwright JB, Schofield C, Conway M, Phillips D, Martin-Silverstone E, Brodrick EA, et al. Multiple axes of visual system diversity in Ithomiini, an ecologically diverse tribe of mimetic butterflies. Journal of Experimental Biology. 2023 Dec 8;226(24):jeb246423.

51. Wright DS, Manel AN, Guachamin-Rosero M, Chamba-Vaca P, Bacquet CN, Merrill RM. Quantifying visual acuity in Heliconius butterflies. Biology Letters. 2023 Dec 13;19(12):20230476.

52. Schindelin J, Arganda-Carreras I, Frise E, Kaynig V, Longair M, Pietzsch T, et al. Fiji: an open-source platform for biological-image analysis. Nat Methods. 2012 Jul;9(7):676–82.

53. Currea JP, Sondhi Y, Kawahara AY, Theobald J. Measuring compound eye optics with microscope and microCT images. Commun Biol. 2023 Mar 7;6(1):1–12.

54. Posit team. RStudio: Integrated Development Environment for R. Posit Software, PBC, Boston, MA. URL http://www.posit.co/. [Internet]. PBC, Boston, MA: Posit Software, PBC; 2023. Available from: http://www.posit.co/

55. Land MF. Visual Acuity in Insects. Annual Review of Entomology. 1997;42(1):147–77.

56. Land MF. Variations in the Structure and Design of Compound Eyes. In: Stavenga DG, Hardie RC, editors. Facets of Vision. Berlin, Heidelberg: Springer; 1989. p. 90–111.

57. Caves EM, Johnsen S. AcuityView: An r package for portraying the effects of visual acuity on scenes observed by an animal. Methods in Ecology and Evolution. 2018;9(3):793–7.

58. Montgomery SH, Merrill RM, Ott SR. Brain composition in Heliconius butterflies, posteclosion growth and experience-dependent neuropil plasticity. Journal of Comparative Neurology. 2016;524(9):1747–69.

59. Lenth RV. emmeans: Estimated Marginal Means, aka Least-Squares Means. R package version 1.8.6, https://CRAN.R-project.org/package=emmeans [Internet]. 2023. Available from: https://CRAN.R-project.org/package=emmeans

60. Warton DI, Duursma RA, Falster DS, Taskinen S. smatr 3– an R package for estimation and inference about allometric lines. Methods in Ecology and Evolution. 2012;3(2):257– 9.

61. Ito K, Shinomiya K, Ito M, Armstrong JD, Boyan G, Hartenstein V, et al. A Systematic Nomenclature for the Insect Brain. Neuron. 2014 Feb 19;81(4):755–65.

62. Habenstein J, Amini E, Grübel K, el Jundi B, Rössler W. The brain of Cataglyphis ants: Neuronal organization and visual projections. Journal of Comparative Neurology. 2020;528(18):3479–506.

63. Garner D, Kind E, Lai JYH, Nern A, Zhao A, Houghton L, et al. Connectomic reconstruction predicts visual features used for navigation. Nature. 2024 Oct;634(8032):181–90.

64. Mota T, Yamagata N, Giurfa M, Gronenberg W, Sandoz JC. Neural organization and visual processing in the anterior optic tubercle of the honeybee brain. J Neurosci. 2011 Aug 10;31(32):11443–56.

65. Rø H, Müller D, Mustaparta H. Anatomical organization of antennal lobe projection neurons in the moth Heliothis virescens. Journal of Comparative Neurology. 2007;500(4):658–75.

66. Farnworth MS, Toh YP, Loupasaki T, Hodge EA, Jundi B el, Montgomery SH. Distinct evolutionary trajectories of two integration centres, the central complex and mushroom bodies, across Heliconiini butterflies [Internet]. bioRxiv; 2025 [cited 2025 Jul 2]. p. 2025.05.02.651904. Available from: https://www.biorxiv.org/content/10.1101/2025.05.02.651904v1

67. Kittelmann M, McGregor AP. Looking across the gap: Understanding the evolution of eyes and vision among insects. BioEssays. 2024;46(5):2300240.

68. Perl CD, Niven JE. Differential scaling within an insect compound eye. Biology Letters. 2016 Mar;12(3):20160042.

69. Rutowski RL, Gislén L, Warrant EJ. Visual acuity and sensitivity increase allometrically with body size in butterflies. Arthropod Structure & Development. 2009 Mar 1;38(2):91– 100.

70. Swihart SL. Acceptance angles of butterfly ommatidia. Journal of Insect Physiology. 1974 Jun 1;20(6):1027–36.

71. Gonzalez-Bellido PT, Wardill TJ, Juusola M. Compound eyes and retinal information processing in miniature dipteran species match their specific ecological demands. Proceedings of the National Academy of Sciences. 2011 Mar 8;108(10):4224–9.

72. Borst A. Drosophila’s View on Insect Vision. Current Biology. 2009 Jan 13;19(1):R36–47.

73. Kinoshita M, Stewart FJ. Cortical-like colour-encoding neurons in the mushroom body of a butterfly. Current Biology. 2022 Feb 7;32(3):R114–5.

74. Galizia CG, Rössler W. Parallel Olfactory Systems in Insects: Anatomy and Function. Annual Review of Entomology. 2010 Jan 7;55(Volume 55, 2010):399–420.

75. Ardin P, Peng F, Mangan M, Lagogiannis K, Webb B. Using an Insect Mushroom Body Circuit to Encode Route Memory in Complex Natural Environments. PLOS Computational Biology. 2016 Feb 11;12(2):e1004683.

76. Gronenberg W. Subdivisions of hymenopteran mushroom body calyces by their afferent supply. Journal of Comparative Neurology. 2001;435(4):474–89.

77. Nishino H, Iwasaki M, Yasuyama K, Hongo H, Watanabe H, Mizunami M. Visual and olfactory input segregation in the mushroom body calyces in a basal neopteran, the American cockroach. Arthropod Structure & Development. 2012 Jan 1;41(1):3–16.

78. Ehmer B, Gronenberg W. Segregation of visual input to the mushroom bodies in the honeybee (Apis mellifera). Journal of Comparative Neurology. 2002;451(4):362–73.

79. Farris SM. Evolution of insect mushroom bodies: old clues, new insights. Arthropod Structure & Development. 2005 Jul 1;34(3):211–34.

80. Mobbs PG. Neural networks in the mushroom bodies of the honeybee. Journal of Insect Physiology. 1984 Jan 1;30(1):43–58.

81. Mega NO, de Araújo AM. Analysis of the mating behavior and some possible causes of male copulatory success in Dryas iulia alcionea (Lepidoptera, Nymphalidae, Heliconiinae). J Ethol. 2010 Jan 1;28(1):123–32.

82. Chakraborty M, Lara AG, Dang A, McCulloch KJ, Rainbow D, Carter D, et al. Sex-linked gene traffic underlies the acquisition of sexually dimorphic UV color vision in Heliconius butterflies. Proceedings of the National Academy of Sciences. 2023 Aug 15;120(33):e2301411120.

83. McCulloch KJ, Yuan F, Zhen Y, Aardema ML, Smith G, Llorente-Bousquets J, et al. Sexual Dimorphism and Retinal Mosaic Diversification following the Evolution of a Violet Receptor in Butterflies. Molecular Biology and Evolution. 2017 Sep 1;34(9):2271–84.

