## Supplementary Information for "Conservation of sensory pathways implies a localised change in the mushroom bodies is associated with cognitive evolution in *Heliconius butterflies*"


**Authors:**

Elizabeth A. Hodge[a,b,*], Denise D. Dell'Aglio[a,b], Antoine Couto[c], W. Owen McMillan[b], Max S. Farnworth[a,†], Stephen H. Montgomery[a,b,†,*]

**Affiliations:**

[a] Evolution of Brains and Behaviour lab, School of Biological Sciences, University of Bristol, Bristol, United Kingdom

[b] Smithsonian Tropical Research Institute, Gamboa, Panama

[c] Evolution Genomes Behavior and Ecology, IDEEV, Université Paris-Saclay, Gif-sur-Yvette, France

### Supplementary Figures

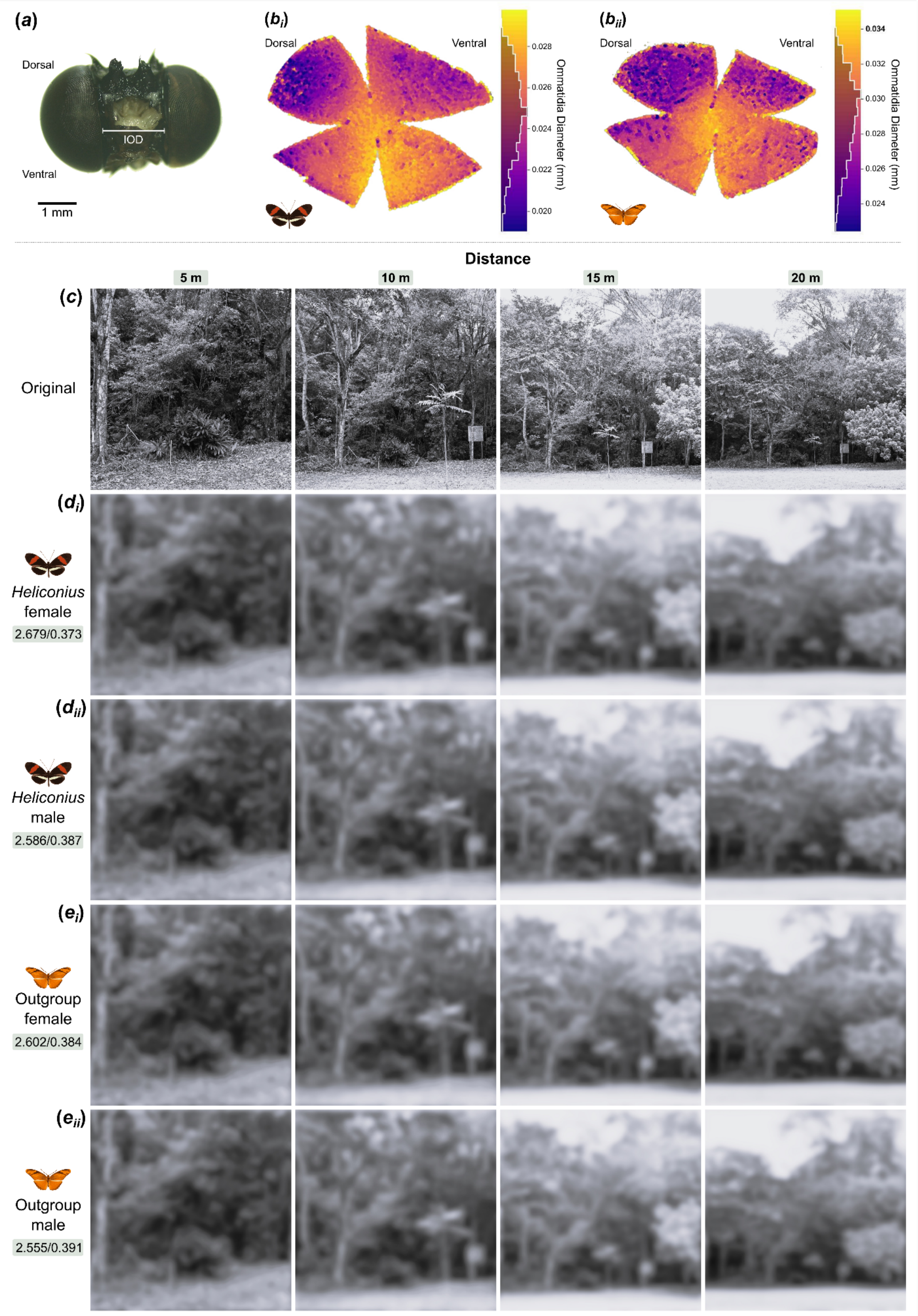


**Figure S1. Visual acuity at different distances across Heliconiini butterflies.** Eye morphology and visual acuity across *Heliconius* and outgroup Heliconiini. **(a)** Front view head photograph example of *Heliconius erato demophoon* with the interocular distance (IOD) measurement indicated. **$b_i$/$b_{ii}$** Example outputs from the ommatidia detecting algorithm of a *Heliconius* (**$b_i$**) and outgroup Heliconiini species (**$b_{ii}$**). Points indicate the different ommatidia detected by the program, with colour denoting diameter of the individual ommatidia (mm). **(c-e)** Morphological visual acuity in male and female Heliconiini demonstrating ability to distinguish main canopy features at distances of 5 m, 10 m, 15 m and 20 m. Visualisation of the scenery were generated with the *AcuityView* package in R from photos the photos in **c**. Values show morphological estimates of the minimum resolvable angle in degree/ acuity in cycles-per-degree.

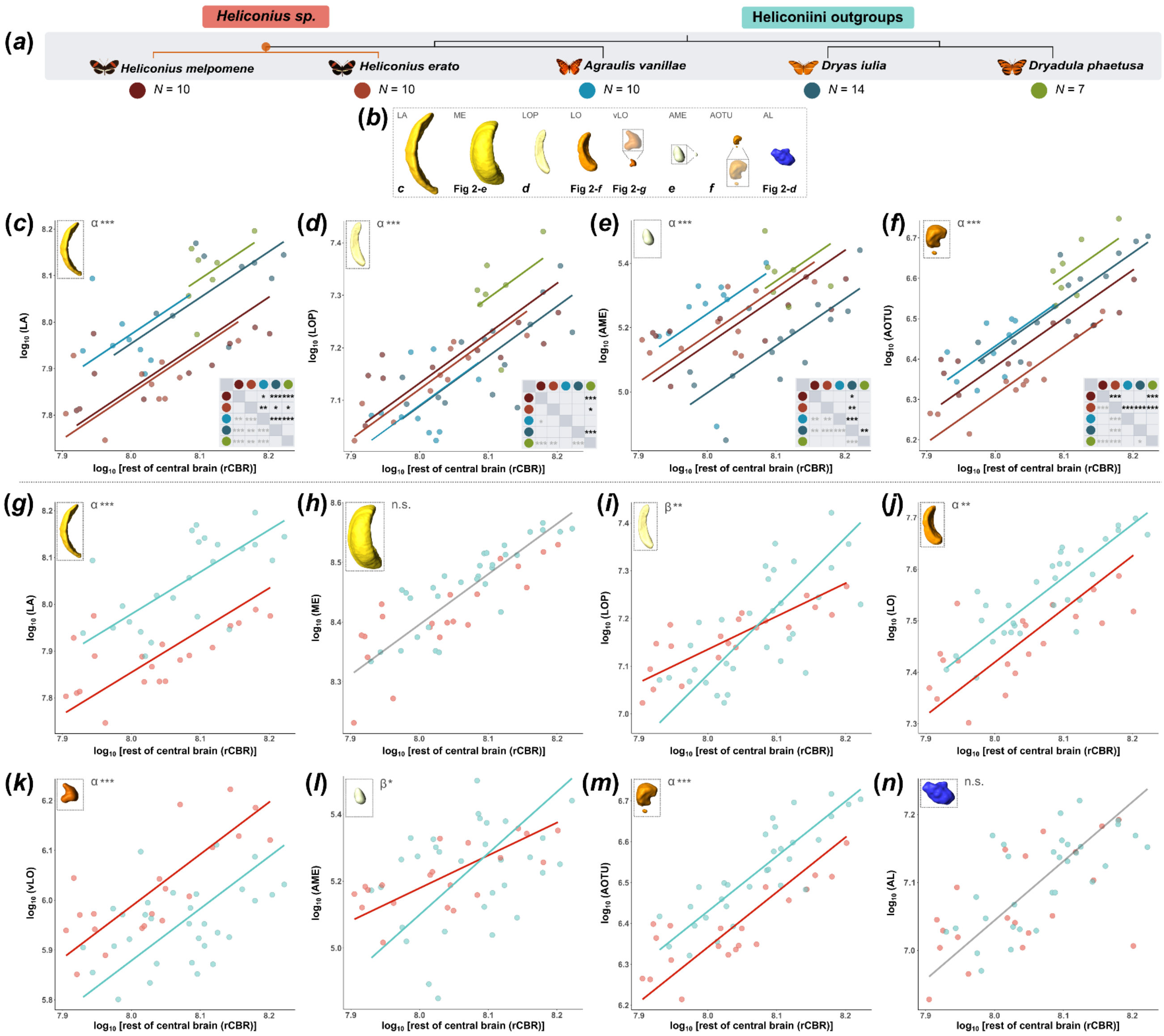


**Figure S2. Sensory neuropil investment across Heliconiini species. (a)** Heliconiini phylogeny with evolution of pollen feeding in *Heliconius* indicated in orange. Heliconiini outgroups are blues and greens, with *Heliconius* species in reds. **(b)** Surface reconstructions of the sensory neuropils moving from the periphery (left) to central brain (right). **(c-f)** Scaling relationships between four of the sensory neuropils (LA, LOP, AME and AOTU) and the rest of the central brain (rCBR) between the five Heliconiini species. Red: *Heliconius* species, blue/green: non-pollen feeding Heliconiini. Matrix of pairwise species difference. Level of significance indicated before (light grey) and after p-adjustment (black). **(g-n)** Heliconiini outgroup versus *Heliconius* scaling relationships of the eight neuropils with the rCBR. A single grey line is presented where no group differences in the scaling components were found. Type of scaling relationship (β/slope-shift, α/grade-shift, major axis shift) is indicated in the top left corner of each structure plot. (*p < 0.05, **p < 0.01, ***p < 0.001).

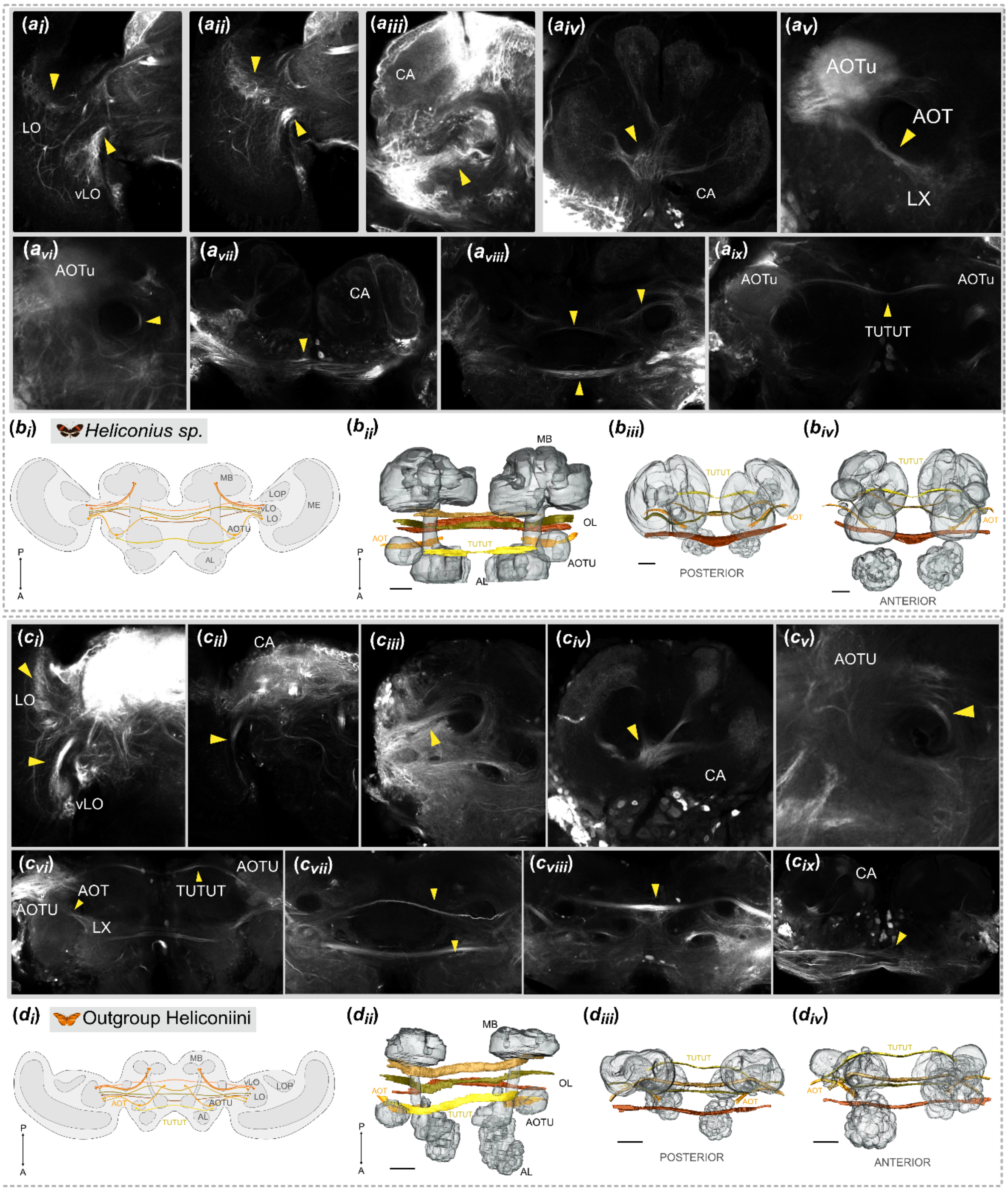


**Figure S3. Major visual pathways in Heliconiini butterflies.** Visual tracts and commissures in *Heliconius* **(a/b)** and outgroup Heliconiini **(c/d)**. Prominent tracts are indicated by the yellow arrows. Visual calyx input in the outgroup Heliconiini (*D. iulia and E. isabella*) and *Heliconius* (*H. melpomene* and *H. hortense*) is identified through calyx lobe injection and optic lobe injection showing input originating from the lobula (LOB) and ventral lobula (vLO) **a/c$_{\mathbf{i-iv}}$**. Optic inputs to the AOTU and lateral complex along the AOT identified through optic lobe injection and GAD-Tubulin staining **c$_{\mathbf{v}}$**. Four optic commissures in *E. Isabella* identified via optic lobe injection and *H. hortense* **c$_{\mathbf{vi-ix}}$**. Schematic of all of the major visual tracts in *Heliconius* **b$_{\mathbf{i}}$** and outgroup Heliconiini **d$_{\mathbf{i}}$**. 3D segmentations of the anterior optic tract (AOT) and tubercle to tubercle tract (TUTUT) as well as three of the four major visual commissures based on staining from a representative *H. melpomene* **b$_{\mathbf{ii/iv}}$** and *D. iulia* **d$_{\mathbf{ii/iv}}$** from a dorsal-ventral and anterior-posterior perspectives. The optic lobes were not available in the scans but their location is indicated in the structural labels in **b$_{\mathbf{ii}}$/d$_{\mathbf{ii}}$** (OL). Scale bars represent 100 µm.

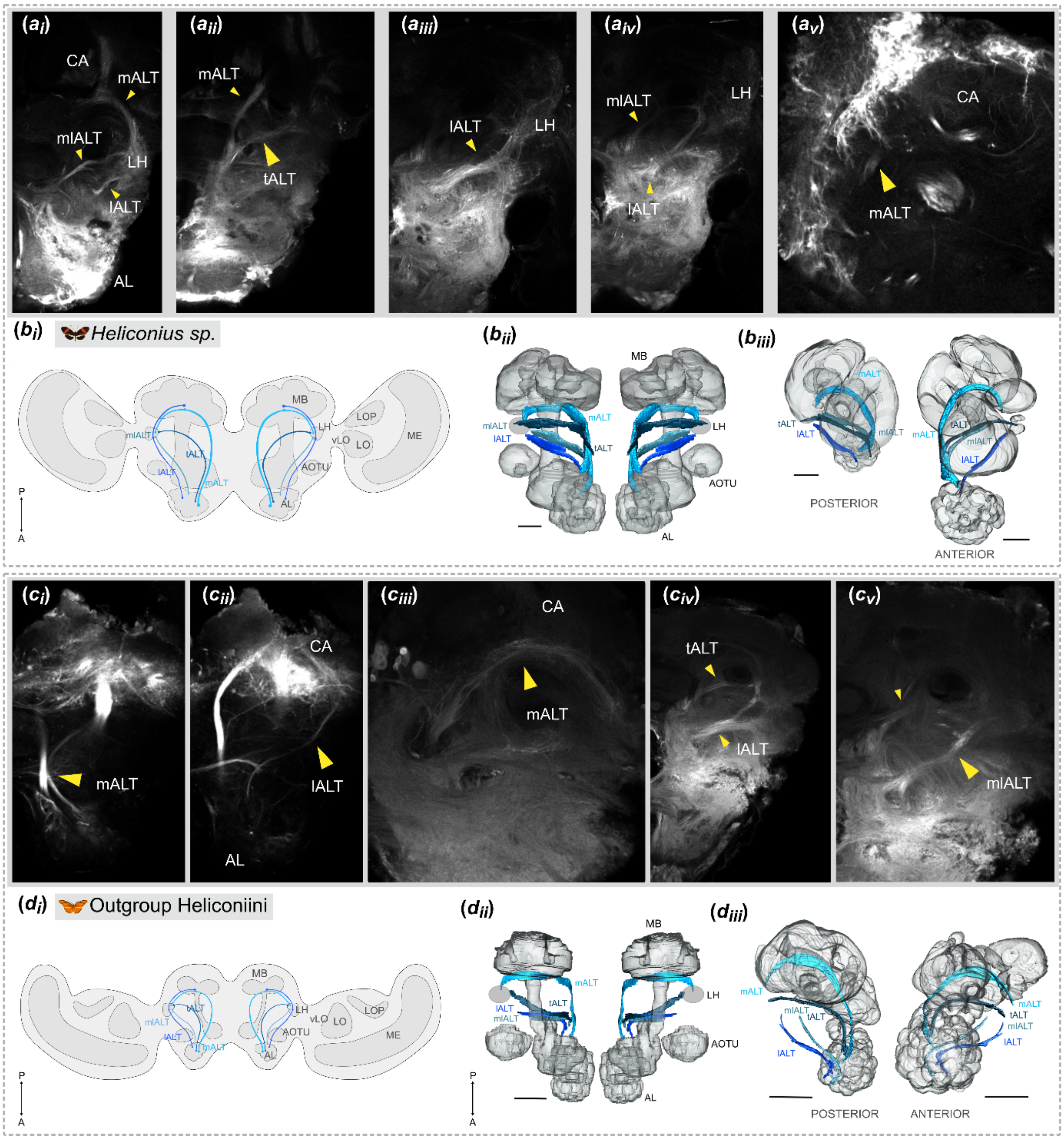


**Figure S4. Major olfactory pathways in Heliconiini butterflies.** Olfactory tracts in *Heliconius* **(a/b)** and outgroup Heliconiini **(c/d)**. Prominent tracts are indicated by the yellow arrows. Tracts are identified based on a combination of antennal lobe and calyx injections across. Olfactory input to the mushroom bodies via the mALT and lALT, as well as input to the LH from the mlALT and tALT represented by AL injections in *H. hortense* $\mathbf{a_i}$-$\mathbf{a_v}$ and *D. iulia* $\mathbf{c_i}$-$\mathbf{c_v}$. Olfactory input to the mushroom bodies via the mALT and lALT, as well as input to the LH from the mlALT and tALT represented by AL injections in *E. Isabella, D. iulia* and *H. hortense.* Schematic of all the major olfactory tracts in *Heliconius* $\mathbf{b_i}$ and outgroup Heliconiini $\mathbf{d_i}$. Schematic of all the major visual tracts in *Heliconius* $\mathbf{b_i}$ and outgroup Heliconiini $\mathbf{d_i}$. 3D segmentations of the mALT, tALT, mlALT and lALT based on staining from a representative *H. melpomene* $\mathbf{b_{ii/iv}}$ and *D. iulia* $\mathbf{d_{ii/iv}}$ from a dorsal-ventral and anterior-posterior perspectives. The lateral horn is indicated as a circle as it was not possible to consistently segment this structure from the available stains. Scale bars represent 100 μm.

### Supplementary Results

#### S1: Differences in neuropil volume across Heliconiini species

Here, we provide more detailed comparisons between the species. Overall, neuropil volume and scaling are largely conserved across the Heliconiini species. In the majority of the neuropils, volumes were conserved between *D. phaetusa* and *D. iulia,* with differences between these species only seen in the accessory medulla and lobula plate, where *D. phaetusa* was larger (Table S2b). The other Heliconiini outgroup species *A. vanillae* was closer to an intermediate between the *Heliconius* species and *D. phaetusa* and *D. iulia.* In only the ventral lobula was *A. vanillae* larger than the other outgroup species and *A. vanillae* and *H. melpomene* were consistently conserved across all of the sensory neuropils (Table S2b). *H. erato* was also conserved with *A. vanillae* in the medulla, accessory medulla, ventral lobula and lobula plate. In most cases, however, *H. erato* had the smallest sensory neuropils of the five species including in comparison to *H. melpomene* (in the medulla, lobula and AOTU) (Table S2b). In the ventral lobula, *A. vanillae, H. melpomene* and *H. erato* were all larger than the outgroup species *D. phaetusa* and *D. iulia.*

The R package *smatr v 3.4-8* (1) was used to more closely assess species-specific allometric scaling relationships between neuropil volumes and rCBR. These analyses revealed interspecific differences in the neuropils, but no increases in investment that were specific to *Heliconius* species following p-adjustment with Šidák correction for multiple comparisons (Figure 2, S2). Indeed, as in the linear model analyses and in contrast to predictions, *Heliconius* species had no greater investment overall than outgroup Heliconiini species. When comparing the two groups, both β shifts in regression slope (accessory medulla and lobula plate) and grade shifts in the intercept (lamina, lobula, ventral lobula and AOTU) were present (Figure 2, Figure S2G-N). These β shifts in the accessory medulla and the lobula plate could indicate a greater rate of sensory neuropil expansion in outgroup Heliconiini than *Heliconius*. However, these β shift patterns are not reflected in the interspecific patterns, where grade shift differences between species are instead present (Table S3ai; Figure S2E), with *D. iulia* having less investment in the accessory medulla than all other species (Table S3aii). Interspecific grade shifts were also found in the lobula plate despite the group β shift, with *D. phaetusa* showing greater investment than the other four species in this structure (Table S3aii; Figure S2D).

In addition to this, the lamina, lobula and AOTU were all larger in outgroup Heliconiini species (Table S3bi, Figure 2, Figure S2G-N). In these neuropils, the between-group grade shift differences are largely reflected in species differences (Table S3aii; Figure S2). In the lamina, differences are seen between *A. vanillae* and the other outgroup Heliconiini species, but scaling is conserved between *D. iulia* and *D. phaetusa*, and between *H. melpomene* and *H. erato* (Table S3aii; Figure S2C). The outgroup Heliconiini species and *Heliconius* species had conserved scaling relationships within their respective groups for the lobula (Figure 2E, Figure S2J). A similar pattern was also observed in the AOTU, however, *H. erato* had lower investment than the other *Heliconius* species, *H. melpomene.* In none of these three sensory neuropils were *Heliconius* species larger than the outgroup Heliconiini species (Table S3aii, Figure S2). In contrast to this trend, in the ventral lobula, the *Heliconius* group were larger than the outgroup Heliconiini species (Table S3bi, Figure 2F, Figure S2K). Both *H. erato* and *H. melpomene* were larger than *D. iulia* and *D. phaetusa.* However, the other outgroup Heliconiini species, *A. vanillae*, was conserved with both *Heliconius* species, and was also larger than the other outgroup species suggesting the ventral lobula expansion is not isolated to *Heliconius* species (Figure 2F).

In the medulla and antennal lobe, group differences did not explain variation in scaling patterns, with interspecific differences found within the *Heliconius* and outgroup Heliconiini groups (Table S3aii, S3bi, Figure S2H/N). A significant grade shift was found between the *Heliconius* species and outgroup species *D. iulia.* With *D. iulia* also having greater investment than *A. vanillae* and differences found between *H. erato* and *H. melpomene.* In the antennal lobe, as expected from the linear model analysis, very minimal species differences were found (Table S3aiii; Figure 2G). A major axis shift along a common slope was seen between *A. vanillae* and *D. phaetusa* with all other species conserved with one another (Figure 2G).

**S2: Sensory projection pathways to the central brain peripheral to the mushroom bodies**

In addition to identifying the main sensory projection pathways to the mushroom bodies, we also provide more detailed descriptions of the identified major optic commissures and additional visual and olfactory tracts to the neuropils within the central brain. The visual and olfactory projections are largely conserved with that seen in other insects, with information integrated across a common set of neuropils with consistent tracts and commissures.

As well as the two major visual tracts going to the mushroom bodies originating in the ventral lobula and lobula (FigureS3A/Ci-iv) and the AOT (Figure S3Av/Cvi) in all of the species, we identified an additional projection connected to the AOTU but the intensity of the staining prevented identification of whether this tract was projecting from or to the AOTU (Figure S3Avi/Cv). This tract projects medially above the peduncle and then turns, projecting laterally back towards the ventrolateral brain region and the optic lobes. We found a conserved pattern of at least four major commissures connect the optic lobes with their contralateral counterpart in Heliconiini (Figure S3Avii-viii/Cvii-Cix). The mushroom body calyces are much larger in the *Heliconius*, resulting in some differences between *Heliconius* and outgroup species in the proximity of the pathways to one another. Nevertheless, the pattern of the projections are the same across the Heliconiini studied, exhibiting consistent anterior-posterior order of the commissures (Figure S3). The most anterior commissure passes below the central complex through the suboesophageal brain region. The second most anterior commissure is found more ventrally than the first, connecting ventral regions of the lobula with the contralateral lobula and passing above the peduncles. The third commissure passes above the central complex but below the peduncles. The most posterior and thickest of the four commissures is composed of many thin fibres connecting the optic lobes passing below the mushroom body calyces (Figure S3Avii/Cix).

With regards to the olfactory projections, the mALT is described across insect species in Coleoptera, Lepidoptera, Diptera, Orthoptera, Blattodea and Hymenoptera (2–10). Input to the calyces from the lALT on the other hand, is less frequently described (6,11). The lALT's projections mainly terminate in the lateral horn, however, injection into the calyces revealed thin fibres that also project to the mushroom bodies (Figure S4Aiii/Cii/Civ). A similar lALT is also observed in dung beetles, though, overall, the tract appears much thicker in Lepidoptera (7). The tALT starts off as part of the mALT and splits from it as the mALT projects posteriorly. Following the split from the mALT, the tALT projects laterally to the lateral horn (Figure S4Aii/Civ). The mlALT splits from the mALT earlier and more anteriorly than the tALT, before also projecting laterally to the lateral horn (Figure S4Ai/Aiv/Cv). The tALT projects more dorsally to the lateral horn than the mlALT. Although the same pattern of olfactory projections was seen across the Heliconiini species, in *Heliconius* the volume of the calyces caused some degree of rotation of the olfactory projections resulting in differences in proximity and rotation anterior-posteriorly (Figure S4B/Dii-iii).

**Bibliography**


1. Warton DI, Duursma RA, Falster DS, Taskinen S. smatr 3– an R package for estimation and inference about allometric lines. Methods in Ecology and Evolution. 2012;3(2):257–9.

2. Farnworth MS, Montgomery SH. Complexity of biological scaling suggests an absence of systematic trade-offs between sensory modalities in Drosophila. Nat Commun. 2022 May 26;13(1):2944.

3. Galizia CG, Rössler W. Parallel Olfactory Systems in Insects: Anatomy and Function. Annual Review of Entomology. 2010 Jan 7;55(Volume 55, 2010):399–420.

4. Habenstein J, Amini E, Grübel K, el Jundi B, Rössler W. The brain of Cataglyphis ants: Neuronal organization and visual projections. Journal of Comparative Neurology. 2020;528(18):3479–506.

5. Homberg U, Montague RA, Hildebrand JG. Anatomy of antenno-cerebral pathways in the brain of the sphinx moth Manduca sexta. Cell Tissue Res. 1988 Nov 1;254(2):255–81.

6. Ian E, Berg A, Lillevoll SC, Berg BG. Antennal-lobe tracts in the noctuid moth, Heliothis virescens: new anatomical findings. Cell Tissue Res. 2016 Oct 1;366(1):23–35.

7. Immonen EV, Dacke M, Heinze S, el Jundi B. Anatomical organization of the brain of a diurnal and a nocturnal dung beetle. Journal of Comparative Neurology. 2017;525(8):1879–908.

8. Ito K, Shinomiya K, Ito M, Armstrong JD, Boyan G, Hartenstein V, et al. A Systematic Nomenclature for the Insect Brain. Neuron. 2014 Feb 19;81(4):755–65.

9. Kanzaki R, Soo K, Seki Y, Wada S. Projections to Higher Olfactory Centers from Subdivisions of the Antennal Lobe Macroglomerular Complex of the Male Silkmoth. Chemical Senses. 2003 Feb 1;28(2):113–30.

10. Kinoshita M, Shimohigasshi M, Tominaga Y, Arikawa K, Homberg U. Topographically distinct visual and olfactory inputs to the mushroom body in the Swallowtail butterfly, Papilio xuthus. Journal of Comparative Neurology. 2015;523(1):162–82.

11. Rø H, Müller D, Mustaparta H. Anatomical organization of antennal lobe projection neurons in the moth Heliothis virescens. Journal of Comparative Neurology. 2007;500(4):658–75.
